# The effects of nitrogen deposition, grazing and land use on Danish acidic grasslands

**DOI:** 10.1101/2022.06.21.493892

**Authors:** Winnie Heltborg, Søren Ranum Wendtland Danielsen, Marianne Haugaard-Christensen, Christian Frølund Damgaard

## Abstract

It has been hypothesized that a strong interplay of land-use history may explain the weakening of the response of species richness to increased levels of nitrogen deposition in acid grasslands. In this study, we use an interpretation and grouping of crop codes from the farmers Field and Fertilizer Plan as an indicator for land use to test this hypothesis. We find that land-use, significantly, affects several indicators of the conservation status of Danish acid grasslands. The study clearly concludes that present and historical management and land-use is important for understanding the differences and changes in the conservation status of acid grassland.

## Introduction

A strong negative correlation between species richness of acid grasslands and atmospheric nitrogen deposition has been demonstrated in the UK (Stevens 2004) and on a larger spatial scale across the Atlantic biogeographic Zone of Europe (Stevens *et al*. 2010). When combining data from the European study with a large data set from a Danish monitoring program, only a non-significant decrease in species richness with nitrogen deposition was found at the Danish sites (Damgaard *et al*. 2011). This negative result may partly be due to a relative small range in nitrogen deposition at the Danish sites, but it was also hypothesized that the observed species richness could be affected by other interfering factors such as present and historical management and land-use. Furthermore, the Danish botanical definition of acid grassland is somewhat broader than the definition used in the study by Stevens *et al*. 2010. The Danish definition includes naturally species poor grasslands, grass heathland and acidic grasslands of less favourable conservation status, and it was suggested that the responses in species diversity in the Danish study could be affected by this broader definition (Damgaard *et al*. 2011).

I this study we combine data on conservation status, nitrogen deposition, grazing and land use, to test the hypothesis that land-use mediate the effect of nitrogen deposition in acid grasslands, and to re-evaluate the significance of the effect of nitrogen deposition on several response indicators when including interactions of these other factors.

Acid grassland is, under Danish conditions, a semi-natural habitat type and persists only through continuous disturbance by livestock grazing or cutting [ref]. A large majority of the acid grasslands in Denmark is owned and/or managed by farmers. As part of these farmers apply for single payment or agri-environmental schemes they are obligated to deliver a “Field and Fertilizer Plan” for each field on the farm with information on the different types of crops, including different types of permanent grass and set-aside land. In our study we use this reported information as a proxy for land-use to evaluate the effects of land use on the conservation status of acid grasslands.

Due to the intensification of agriculture, farming by grazing is no longer profitable and abandonment is a threat to the conservation of grasslands (Timmermann *et al*. 2014). To evaluate the importance of grazing management on the conservation status of acid grassland, we investigate the effect of grazing by livestock on the conservation status, as well as the possible effect of the type of grazing animals.

## Materials and Methods

### Acid grasslands sites

Danish acid grassland sites have been monitored since 2004 as part of the Danish National habitat monitoring program, NOVANA (Nielsen et al. 2012). The vegetation structure is recorded using a random stratified sampling process in 20, 40 or 60 plots per site, depending on its size. At each plot, vegetation data of vascular plants is recorded by pin-point measure (n = 16) in a 0.25 m^2^ quadrate and a complete species list in a 78.5 m^2^ circle (radius = 5 m) centered on the sample quadrate. Additionally, a number of physio-chemical soil properties, vegetation structural properties, and other ecological indicators were measured (Nielsen et al. 2012).

The vegetation types of all plots were classified according to the habitat classification system used in the European Habitat Directive (EU 2003) and only plots that were classified as acid grassland (EU habitat type 6230; species-rich *Nardus* grasslands, (Ejrnæs et al. 2004) and monitored after 2007 were used in the analysis. Some plots were monitored several times, in those cases only the last monitoring event was used in the analysis. A total of 2262 plots from 103 sites were used in the analysis.

### Ecological indicators

The conservation status of the plant communities in the acid grasslands were measured by a number of ecological indicators (Table 1). We choose indicators form the Danish monitoring program representing effects on different functional and structural features of acidic grasslands (Nielsen et al. 2012). These indicators are amongst others used to evaluate the current status of the acid grasslands, and are expected to respond to changing levels of nitrogen deposition, intensity of land use and grazing management (Table 1). Indicators for vegetation height and tree/shrub cover should also help evaluate whether crop codes and our grouping and ranking of these can be used as a valid indicator for land use.

**Table 1.** Ecological indicators, and their features.

| Indicator |  |
| --- | --- |
| Species, number of species per sample circle ($r = 5 \text{ m}^2$) | The number of species per quadrat in each plot was used as a simple indicator for species diversity. A decline in species diversity as a result of increased levels of nitrogen deposition has been demonstrated in several studies (kilde) and a high species diversity is usually believed to be associated with good ecological status of the acid grasslands. Species diversity could also be affected by grazing and land use as cutting or fertilizing |
| Indicator species, number of species per sample circle ($r = 5 \text{ m}^2$) | In Denmark, there is no official lists of typical species for the annex 1 habitat types. The Danish monitoring programme includes expert made lists of "indicator species". These species are chosen for their strength to indicate even relatively low levels of impacts negatively affecting the natural habitats, yet having a relatively high occurrence and representing the natural variety within habitat types (ref). Indicator species chosen for their sensitivity for disturbance or threats should be expected to respond to changing levels of deposition, land use and grazing regime. A high number of indicator species is an indication of continuity and favourable management and is usually believed to be associated with good ecological status of the acid grasslands. |
| Nutrient ratio, EIIN/EIIR, mean value per sample square ($1 \text{ m}^2$ weighted by frequency, pin-point) | Nutrient ratio is a corrected Ellenberg indicator. The mean Ellenberg N is strongly correlated with mean Ellenbergs R (ref). A high nutrient ratio indicates that number of species with high Ellenberg N, or high nutrient requirements, is overrepresented related to what would be expected consulting the acidity of the area only. Knowing that acidic grasslands in Denmark includes more or less naturally species poor acid grasslands (grass heathlands), this is a highly relevant indicator. |
| Grimes competitive strategy index GCc mean value per sample square ($1 \text{ m}^2$ weighted by frequency, pin-point) | Grimes CSR-system, is a way to group plants in relation to their occurrence along combinations of environmental gradients of disturbance and stress. Plants occurring in a stable, nutrient rich (low stress) environment usually show a competitive strategy (GC) with at high growth rate and ability to outcompete plants showing stress tolerant (GS) or ruderal strategies (GR) in this type of environment (link grimes 1997). The strategies are correlated and plants show different combination of the three strategies. Ejrnæs & Bruun (2000) have calibrated the strategies of plants occurring across dry grasslands by assigning points to each species representing their relative C,S and R strategy into a total of 12 points for the combination of the strategies (Ejrnæs & Bruun 2000). These indices are used in this study. The Grimes competitive strategy, index GCc is correlated to Ellenbergs N indicator and we expect the of GCc to increase at higher levels of nitrogen depositions or fertilizing which can favour nutrient requiring fast growing species. |
| Grimes stress Strategy index, GSc | Plants showing a stress tolerant strategy are favoured in the nutrient poor habitats of acid grasslands. Eutrophication through nutrient enrichment from |
| mean value per sample square (1 m<sup>2</sup> weighted by frequency, pin-point) | nitrogen deposition or fertilizing is expected to favour plants showing a more competitive strategy. Grazing and cutting will usually lower the nutrient status of the grasslands leading to more favourable conditions for stress tolerant species. |
| Vegetation height (height of the lower vegetation layer, forbs grasses ect. 1 m<sup>2</sup>, pin-point) | Nitrogen deposition can favour competitive tall growing species and there is a correlation between GC<sub>C</sub> and the potential height of the vegetation. Cutting and grazing will however have an immediate effect on vegetation height when it is a direct measurement as in the NOVANA program and then it is a more appropriate an indicator of management as land use or grazing. In the study the indicator could also help us evaluating whether the ranging in intensities of land use is appropriate. |
| Tree/shrub cover, (relative cover of trees and shrubs within a circular plot with a 5 m radius) | Some shrub and tree vegetation is natural to acid grasslands, abandonment will lead to an increase in coverage, and nitrogen deposition is believed to accelerate encroachment. Land use, cutting and grazing will however prevent or delay these effects, and tree/shrub cover can be an indicator of abandonment, management as land use or grazing and the intensity of land use. In the study the indicator could also help us evaluating whether the ranking of land use is appropriate. |
| Forb ratio (cover of forbs/grasses in 1 m<sup>2</sup> plots, pin-point) | Forbs are believed to be more sensitive to eutrophication than grasses, a change in the ratio between these lifeforms can indicate changes in the nutrient status of the acid grasslands and a change in vegetation towards less or more favourable conservation status as it directly effects the vegetation. Effect of grazing? |

### Nitrogen deposition

The expected nitrogen deposition at each plot was calculated using an atmospheric deposition model (Ellermann et al. 2007) as the average of the model predictions in 2008 and 2011. It is known that the relative difference in nitrogen deposition between sites is relatively stable across years on a long term average (Matejko et al. 2009) and the model-predicted nitrogen deposition was used as a proxy for the build-up of N in the soil due to atmospheric deposition. The range of the total nitrogen deposition in Danish acid grasslands ranges from 6 – 18 Kg N kg ^-1^ year^-1^.

### Grazing

We used aerial photos from ‘Area information’ to identify the type of grazing animals. ‘Area information’ is a website-based Geographical Information System, GIS, which presents data geographically referred on map layers. Data regarding nature protection, conservation, building lines, agriculture, planning, soil contamination, and groundwater is presented (The Danish Natural Environment Portal). It is also possible to locate grazing animals and determine if they were sheeps og cows on specific areas by using orthophoto map layers and the zoom function on the website.

Coordinates from each individually plot from the NOVANA stations were entered ‘Area information’ and each plot became marked as grazed or not grazed. Subsequently the NOVANA stations and the adjacent areas to the stations were examined for grazing animals in 2008, 2010 and 2012. If grazing animals were present one of the aforementioned years, it was assumed that the grazing animals (and the same kind of grazing animals) were present the years in which the NOVANA stations were selected.

### Land use

Due to the lack of natural constraints to cultivation Denmark is an intensively cultivated land. The Danish definition of acid grassland is adapted to these conditions and includes more species poor grasslands and grassland of less favourable conservation status such as grasslands on ex-arable agricultural improved land (Damgaard *et al*. 2011). Land use is recognized as a major driver og biodiversity and ecosystem functioning (Blütgen et al (2021). But the monitoring program does not include fulfilling records of management practice or land-use history and this hypothesis could not be tested using data from program alone.

A large proportion of the acid grasslands is owned by and/or managed by farmers. The vast majority of the acid grasslands have been protected against regularly ploughing and cropping since 1992. A few has legally been be applied small amounts of fertilizer many years since. Due to regulation and as part of the application for single payment or agri-environmental schemes, all farmers are obligated to hand-in a “Field and Fertilizer Plan”.

The plan includes a classification of each field on the farm. There is a class for every type of crop, including classes of permanent grassland of different yield and mangement and a class for set-aside land. Crops, soil fertility, and for grasslands, also the use of grazing and cutting, provide a base for calculating a nitrate quota on farm level.

Fields of acid grasslands on farm are included in the Field and Fertilizer Plan as permanent grassland or set aside land. The most productive areas can be used for silage, hay or winter fodder productions. Less productive areas are set aside or managed extensively by grazing and/or cutting or at least trimming.

In this study we use an interpretation of classes for the variety of grasslands as an indicator for land use. The classes have been public available since 2008 and has been extracted from The Danish Agricultural Agency. The classes are ranked and grouped in three groups of land use intensity “intensive”, “extensive” based on expert-interpretations of the classes. and “no agricultural use” for unclassified plots. Grasslands with no classification usually does not belong to a farm and fertilizing is very unlikely.

Intensity was ranked by factors as probability of fertilizing, and level and type of disturbance such as cutting intensity, grazing or trimming. No fertilizing and moderate disturbance by grazing was believed to have the most positive effects for the conservation status of acidic grasslands and assigned to the extensive land use group. The resulting groups were aggregated with the data from the monitoring program for the same year, or the following, and included in the statistical analysis. We expect that more intensively used grasslands may have a correlation to the historic land use, as the farming interests often will be higher on more productive areas where yields are higher and these more productive grassland may have a history of fertilizing and tillage (pasture like – rough grazing).

### Statistical analysis

The hierarchical data on conservation status was analysed in a mixed effect model with nitrogen deposition, grazing, and land use as fixed factors and site as a random factor using the R packages *nlme* (Pinheiro et al. 2013) and *lme4* (Bates et al. 2015). The R package *lme4* was used when analysing *species* and *indicator species*, where the variation was assumed to be Poisson distributed, otherwise the residual variation was assumed to be normally distributed and analysed using the R package *nlme*. The distribution of the residuals was checked visually in residual plots. The categorical fixed factors *grazing* and *land use* were analysed using contrasts. Two linear models were used in the statistical analysis:

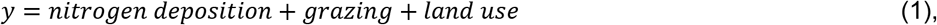

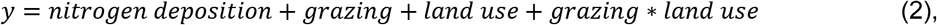

and the model with the lowest AIC was selected as the model that was best supported by the data. In all cases this selection criteria was equal to selecting the simpler model (1) when the interaction term in model (2) was not significant.

## Results

### Nitrogen deposition

The deposition had a significant positive effect on the nutrient ratio (*p= 0.0001*) (table 2), thus showing that nutrient requiring plants have a significant higher occurrence in plots with a higher deposition, related to what would be expected based on the acidity of the area only.

**Table 2.**

| Indicator | Species |  | Indicator species |  | EIIN/EIIR |  | GCc |  | GSc |  | Vegetation height |  | Trees/schrub cover |  | Forb ratio |  |
| --- | --- | --- | --- | --- | --- | --- | --- | --- | --- | --- | --- | --- | --- | --- | --- | --- |
|  | Mixed model: Species ~ Ndep + animals + land use + Land use * animals |  | Mixed model: Indicator species ~ Ndep + animals + land use |  | Mixed model: Nutrient ratio ~ Ndep + animals + land use + Land use * animals |  | Mixed model: GC ~ Ndep + animals + land use |  | Mixed model GS ~ Ndep + animals + land use |  | Mixed model: Vegetation hight ~ Ndep + animals + land use |  | Linear mixed model: Trees and schrubs ~ Ndep + animals + land use + land use * animals |  | Linear mixed model: Herbs ~ Ndep + animals + land use + land use * animals |  |
| Treatment | value | p-value | value | p-value | value | p-value | value | p-value | value | p-value | value | p-value | value | p-value | value | p-value |
| **N dep** | 0,0013 | 0,252 | 0,0061 | 0,7271 | 0,0153 | 0,0001 *** | -0,0387 | 0,0138 * | 0,0665 | 0,0546 | -0,286 | 0,0391 * | -0,155 | 0,5033 | -0,001 | 0,6377 |
| **Ungrazed, grazed** | 0,0013 | 0,878 | 0,0293 | 0,1274 | 0,0016 | 0,8013 | -0,0804 | 0,0280 ** | -0,028 | 0,5795 | -2,148 | 0 *** | -0,985 | 0,0169 * | -0,014 | 0,0042 ** |
| **Sheep, cattle** | 0,1261 | 1E-07 *** | 0,0563 | 0,29 | -0,056 | 0,0006 ** | -0,091 | 0,1612 | -0,165 | 0,1953 | -3,359 | 0 *** | 0,9103 | 0,3801 | 0,0502 | 0,0001 *** |
| **No use, used** | -0,003 | 0,753 | 0,0254 | 0,0958 | -0,032 | 0 *** | -0,0637 | 0,0069 * | -0,064 | 0,128 | -0,707 | 0,002 ** | -4,051 | 0 *** | -0,015 | 0,0167 * |
| **Extensive, intensive** | -0,007 | 0,724 | -0,105 | 0,0001 *** | 0,0172 | 0,1673 | 0,1041 | 0,0065 * | -0,229 | 0,0016 ** | 0,1215 | 0,7342 | 0,3771 | 0,6348 | 0,0036 | 0,7056 |
| **Ungrazed, grazed: no use, used** | -0,006 | 0,244 |  |  | -0,006 | 0,1707 |  |  |  |  |  |  | 0,4548 | 0,1162 | -0,005 | 0,1496 |
| **Sheep, cattle: no use, used** | -0,013 | 0,383 |  |  | 0,0365 | 0,0006 *** |  |  |  |  |  |  | 0,2991 | 0,673 | 0,0031 | 0,7185 |
| **Ungrazed, grazed: extensive, intensive** | 0,0009 | 0,918 |  |  | -3E-04 | 0,9597 |  |  |  |  |  |  | -0,191 | 0,6627 | -4E-04 | 0,942 |
| **Sheep, cattle: extensive, intensive** | -0,01 | 0,684 |  |  | -0,008 | 0,5987 |  |  |  |  |  |  | -1,602 | 0,099 | 0,0104 | 0,3773 |

Significans codes: 0 '\*\*\*\*' 0,001 '\*\*\*' 0,01 '\*\*'
Model 2 was the best model when testing effects om nutrient ratio, EIIN/EIIR, tree cover, species and forb ratio. All combinations of land use and animals were tested.

The deposition has a significant negative effect on GC_c_ (*p = 0.0138*) and vegetation height (p=0.0391).

### Grazing

When contrasting no grazing with grazing, there was no significant effect on species richness, but when contrasting the effects of grazing by sheep and cattle there was a strong significant positive effect of grazing with cattle as opposed to grazing by sheep (*p<0.0000)*. The nutrient ratio was also significantly affected by the type of grazing animals. When contrasting no grazing with grazing, no significant effect of grazing could be seen, however there was a significant difference in the effect of grazing by cattle when contrasting with grazing by sheep (p=0.0006). The two types of animals had significantly different effects on the nutrient ratio (fig??). Grazing by cattle reduces the nutrient ratio relative to grazing by sheep. Grazing has a significant negative effect on vegetation height as opposed to no grazing (*p<0.0000*) and the effect of grazing by cattle is significantly more negative than grazing by sheep (p<0.0000). Grazing also has a significant negative effect on GC_C_ when contrasting no grazing (*p=0.0280*). But no significant effect on GC_C_ was found when contrasting grazing by cattle with grazing by sheep.

Grazing has a significant negative effect on tree/shrub ratio p=0.0169 and on forb ratio p=0.0042. When contrasting the effects of grazing by sheep with cattle there is no effect on tree/shrub cover but a strong significant increase in forb ratio for cattle as opposed to sheep p= 0.0001.

### Land use

There was a significant negative effect of agricultural land use on EIIN/EIIR and GCc, vegetation height, tree/shrub cover when contrasting no use (table??). No agricultural use, was increasing the EIIN/EIIR-ratio. Agricultural land use leads to a small decrease in the forb ratio p= 0.0167 when contrasting with no use. There is a significant decrease in the number of indicator species p= 0.0001 and stress tolerant species p=0.0065 with intensive land use as opposed to extensive land use.

### Interacting effects, grazing and land use

There is a significant effect on nutrient ratio of combinations of agricultural land use and type of grazing animal. The combination of sheep and no agricultural land use has the highest mean value for nutrient ratio. Consistent with the general indication that grazing by cattle in contrast to sheep tend to have the most positive effects on the different indicators.

## Discussion

Using a wider definition of acid grasslands can lead to difficulties in tracking changes in the ecological status by use of indicators, since the response in some of the indicators in the specific area holding the habitat type may be within the variety of the indicators between areas holding the habitat type. Danish acid grasslands include more species-poor varieties of the habitat type and grasslands of less favourable conservation status. Nutrient enrichment may lead to an increase in species diversity for the most species poor areas and a decrease for the most nutrient enriched grasslands resulting in a humpbacked curve for changes in species diversity along the nutrient gradient within the type(ref). Different levels of disturbance can further scatter the responses in species diversity. Ex-arable and fertilized grasslands can contain altered compositions and raised levels of nutrients and the flora will be less sensitive to deposition. Species diversity can also be altered through many years in ex-arable fields due to lack of nearby species pools as these habitat types often are small and fragmented areas in the landscape in Denmark (ref)

Nitrogen deposition has a significant positive effect on EIIN/EIIR ratio, indicating that there is an overrepresentation of nutrient requiring plants with increasing levels of nutrient deposition. The deposition has a significant negative effect on GC_c_ (*p = 0.0138*) and vegetation height (p=0.0391). Nitrogen deposition is expected to favor competitive tall-growing species (ref..)(table).

The negative effect on Grimes Competitive Index (and vegetation height) indicates that though there is an increase in the frequency of nutrient requiring plants with an increase in nitrogen deposition, these changes is not leading to an increase in GC_c_. Other studies have shown an increase in GC_c_ and potential vegetation height resulting from changes in the vegetation and loss of species in semi natural habitats in Denmark throughout the years (Timmermann *et al*. 2014). This is corresponding with the results of the monitoring programme evaluating the status and changes in the acid grasslands where increased levels of GCc is shown from 2004-2009 (reff???).

A decrease of Grimes Competitive Index with increased levels of nitrogen deposition, as shown in this study could be a result of a more intensive use (increased utilization and disturbance) of the most productive areas. That leads to a shift towards species showing ruderal strategies especially since there is no corresponding increase in GSc and the three strategies are directly correlated.

Grazing and agricultural land use both have significant effects on GC_c._ Grazing and land use decreases the GC_c_ and when contrasting the effects of more intense land use with more extensive use there is a positive effect of intensity on GC_c._ This indicates that both abandonment or previous and/or present agricultural improvement and intense use have negative effects on the conservation status of acid grassland.

Grazing by cattle reduces the nutrient ratio, vegetation height and increases forb-ratio when compared with grazing by sheep. This indicates that grazing by cattle have positive effects on the conservation status of grasslands. Grazing has a significant positive effect on species richness.

I this study the grouping in intense land use can be a result of fertilization and/or previous tillage which would lead to an increase in overall nutrient levels whereas raised nitrogen deposition affects only the accessibility of nitrogen. Despite the fact that land use may include applying small amounts of fertilizers, it also includes some annual utilization by cutting and grazing, which keeps the vegetation low and free of shrubs and trees.

When contrasting intensive land use with extensive, there is a significant decrease in the number of indicator species with intensive land use. This corresponds very well to the results showing that intense land use also leads to a decrease in species showing stress tolerance and an increase in competitive species when contrasting with extensive land use.

Raised levels of nutrient status and previous tillage would be likely to have an effect on the indicator species in the Danish monitoring programme. Since these species are chosen for their strength to indicate previous fertilization and/or cultivation and for species adapted to stressful conditions in nutrient leached areas. Reduced number of indicator species could therefore be a result of present or previous fertilization and tillage. And a reduced number of indicator species will be equal to a poorer conservation status in the Danish monitoring programme.

The intensity of the land use affects all the indicators representing changes to the flora except for the species diversity. But as mentioned above, species diversity may not be a very strong indicator for favourable conservation status for the very widely defined Danish acid grasslands.

The combination of sheep and no land use has the highest mean value for the nutrient ratio. The “no use” group of acid grasslands is likely to include areas owned and managed by others bodies than farmers such as governmental agencies. The areas are managed with grazing mostly for the landscape and cultural values, not for agricultural values. We suspect that sheep may be the preferred animal for the most nutrient poor areas managed by grazing, since sheep require less food and cause less damage to the vegetation cover. However, sheep is known to be a more selective grazer and the result indicate that sheep affects the flora negatively when contrasting with cattle.

Land use has significant positive effects on structural indicators such as tree and schrub cover and vegetation hight. These results are supporting our grouping in no use or used. We believe there is a strong correlation between the productivity and the farming interests in acidic grasslands. This is supported by the results showing intense use have more negative effects on the composition of the flora than extensive use. Land use, as defined and ranked in this study, affect several indicators of conservation status. Therefore the use of crop codes could be valuable in understanding response or lack of responses to other abiotic impacts

## Conclusions

The present study cannot confirm that the major driver of changes in Danish acid grassland is increased levels of nitrogen deposition. Deposition has effects but other factors are very important too.

Land use, as defined and ranked in this study, affect several indicators. The results are supporting theory by Damgaard e*t al*. 2011. Present and historical management and land use is important for understanding the differences and changes in the conservation status of Danish acid grassland.

There are several clear differences in the composition of the flora of intensive land use when contrasting with extensive. There are significantly fewer indicator species in areas of intense use and the conservation status of these is likely to be poor.

Grazing, types of grazing, and land use intensity effects several ecological indicators and could play very important roles in the conservation status and changes in conservation status of the acid grasslands.

And in the light of these results it should be recommended to use a much more integrated approach for nature conservation of Danish acid grasslands. Different, or multiple approaches for improving the conservation status must be considered and it should be advised to pay some attention to how you can reverse or remove effects of previous land use.

Crop codes and our grouping and ranking of these into groups of extensive and intensive use could be useful for further studies and for adapting management plans for the specific conditions of the grassland. A narrow management focus on reducing nitrogen deposition will probably not have a very strong effect on the conservation status.

It would also be advised to use indicators for conservations status that can detect changes across the broad definition of the type. A simple focus on species diversity cannot be advised when considering the broad definition of Danish acid grasslands including naturally very species poor grasslands. Here a decrease in species diversity may be a result of either degradation or improvement depending on the starting point.

## Tables and figures

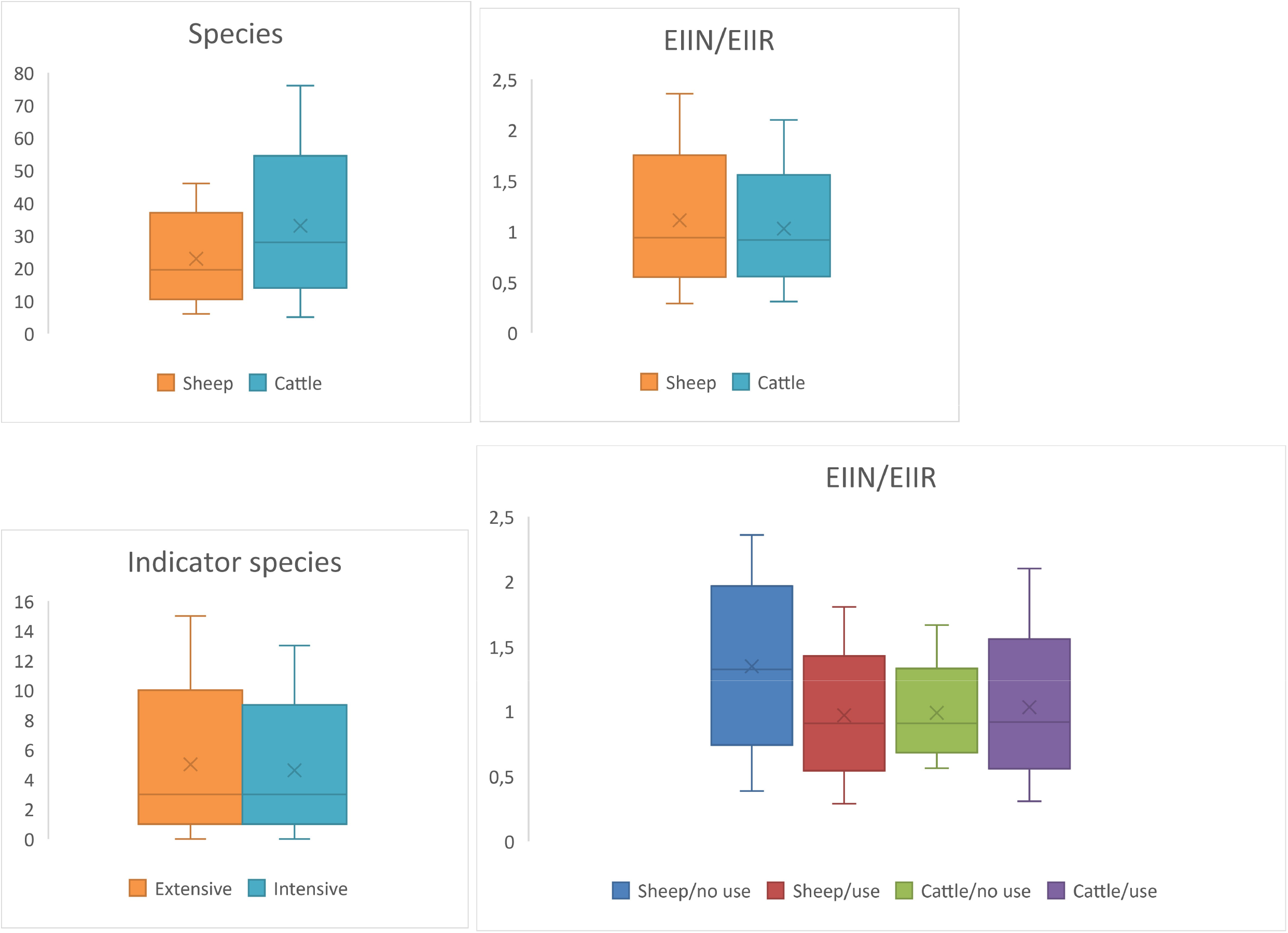

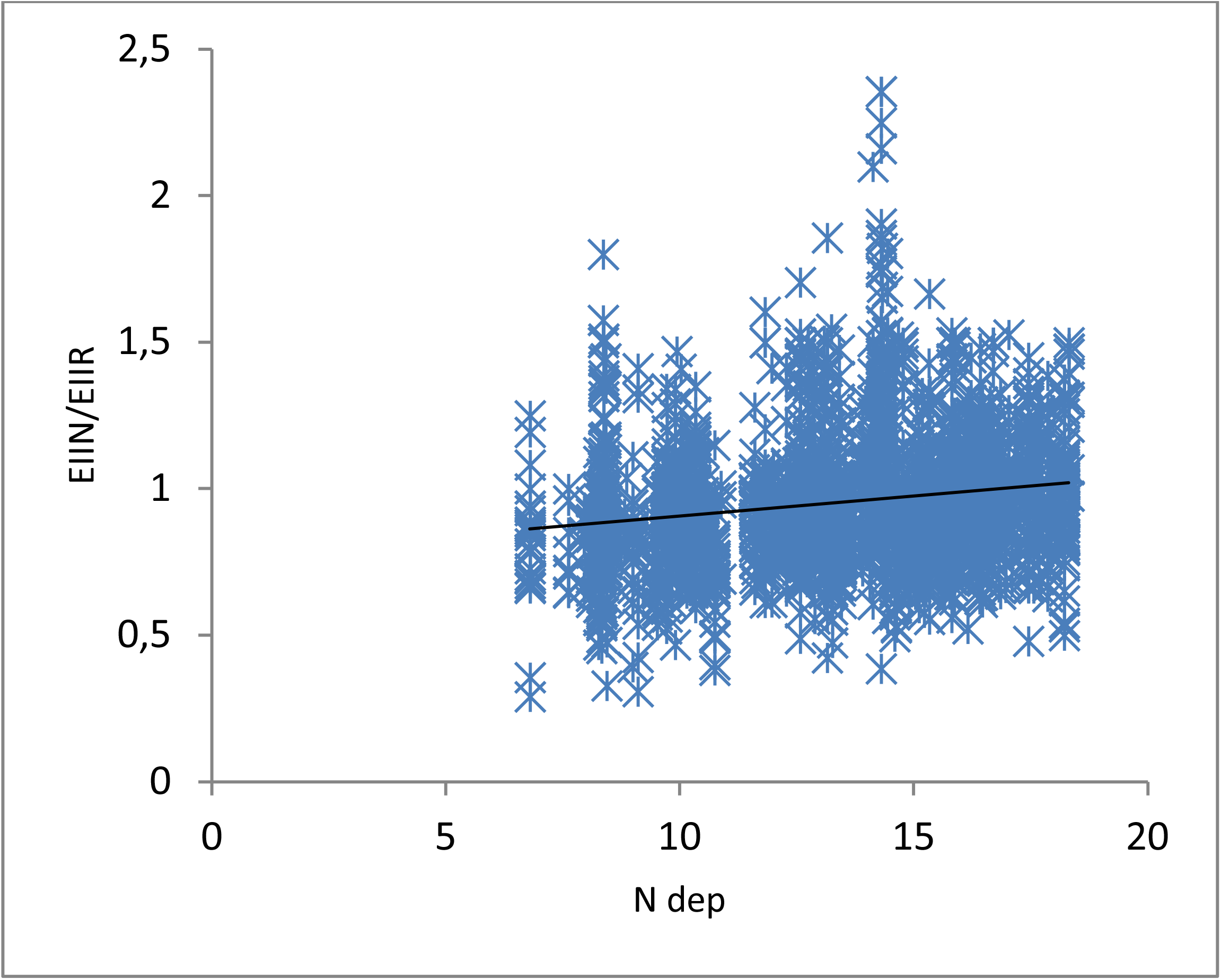

